# Impact of different mutations on Kelch13 protein levels, ART resistance and fitness cost in *Plasmodium falciparum* parasites

**DOI:** 10.1101/2022.05.13.491767

**Authors:** Hannah M. Behrens, Sabine Schmidt, Domitille Peigney, Ricarda Sabitzki, Isabelle Henshall, Jürgen May, Oumou Maïga-Ascofaré, Tobias Spielmann

## Abstract

Reduced susceptibility to ART, the first-line treatment against malaria, is common in South East Asia (SEA). It is caused by point mutations, mostly in *kelch13* (*k13*) but also in other genes, like *upb1*. K13 and its compartment neighbors (KICs), including UBP1, are involved in endocytosis of host cell cytosol. We tested 135 mutations in KICs but none conferred ART resistance. Double mutations of *k13*C580Y with *k13*R539T or *k13*C580Y with *ubp1*R3138H, did also not increase resistance. In contrast, *k13*C580Y parasites subjected to consecutive RSAs did, but the *k13* sequence was not altered. Using isogenic parasites with different *k13* mutations, we found correlations between K13 protein amount, resistance and fitness cost. Titration of K13 and KIC7 indicated that the cellular levels of these proteins determined resistance through the rate of endocytosis. While fitness cost of *k13* mutations correlated with ART resistance, *ubp1*R3138H caused a disproportionately higher fitness cost.

**Significance:** ART resistance is only a partial resistance with a proportion of ring stages surviving physiological ART levels. The correlation of resistance with fitness cost in isogenic lines indicates that fitness cost likely prevents resistance levels permitting survival of all ring stages under physiological ART concentrations. We also found no indication that double mutations in *k13*, including the two most common resistance mutations in SEA, or with non-*k13* genes, are a threat to lead to higher resistance. However, repeated ART exposure increased resistance by mechanisms not altering *k13* gene sequence, indicating changes in the background of these parasites. The disproportionally high fitness cost of *ubp1*R3138H may explain why *kic* mutations affecting resistance are rare and highlights the unique property of K13 to influence endocytosis only in ring stages.

## Introduction

Malaria kills approximately half a million people per year, most of them children in sub-Saharan Africa (*1, 2*). Treatment heavily relies on artemisinin and its derivatives (ARTs) which are typically used in artemisinin-based combination therapies (ACTs) with a partner drug (*3*). More than 10 years ago reduced susceptibility to artemisinin was observed in low malaria incidence settings such as Asia, Oceania and South America. This manifests as delayed parasite clearance after treatment (*3, 4*), caused by some ring stage parasites surviving the ART-pulse, which is brief (a few hours) because of the low half-life of ART drugs (*5*) and can lead to treatment failure (*6*). *In vitro* this reduced susceptibility is measured by the ring stage survival assay (RSA) where survival of >1% of parasites indicates what we call *in vitro* ART resistance (*7*).

The main cause of reduced ART susceptibility are mutations in the *Pfkelch13* gene (*k13*) encoding the K13 protein (*8, 9*), which was recently found to be involved in endocytosis of host cell cytosol in the ring stage of the parasite (*10*). Ten mutations affecting the C-terminal propeller domain of K13 are confirmed to cause reduced susceptibility as evident from delayed clearance in patients (F446I, N458Y, M476I, Y493H, R539T, I543T, P553L, R561H, P574L, C580Y) and others have been associated with reduced susceptibility (*3*). Several other proteins, mostly from the K13-compartment, confer *in vitro* ART resistance in RSA either when downregulated or when mutated (*10–14*). One of them is *ubp1*, for which the R3138H was shown to result in moderate resistance in RSA (*10*) and which was associated with reduced susceptibility in patient samples from Asia, although it occurred only rarely (*15*). The contribution of mutations affecting K13 compartment proteins (other than K13) to reduced ART susceptibility in endemic countries is largely unknown.

Resistant clinical isolates and laboratory parasites carrying the resistance-associated mutations *k13* R539T and *k13* C580Y contain reduced amounts of K13 (*10*, *16–18*). Artificially modulating K13 levels indicated that the reduced K13 levels lead to resistance (*10*, *16*, *19*). It is currently unclear whether other *k13* mutations, like R561H, confer resistance through the same mechanism or affect the functionality of K13.

While ART-resistance mutations are frequent in South East Asia, up to 100% in some locations (*20–22*), they are rare in Africa (*21*, *23*, *24*). Six validated resistance-conferring *k13* mutations (M476I, P553L, R561H, P574L, C580Y and A675V) have occasionally been reported in African countries at moderate to low frequency of 4.1% or less (*25–27*). Two exceptions are the R561H mutation in Rwanda, which was reported more frequently since 2019, most recently up to 16% (*28–30*) and the 11% of A675V mutation in Uganda in 2019 (*27*). It is not clear whether further resistance mutations exist in Africa. Most studies focused on mutations occurring in Asia and there is no systematic *k13* surveillance in Africa. In agreement with the rare occurrence of resistance mutations, delayed parasite clearance occurs at less than 2% in Africa (*31*), with the exception of the Masaka region in Rwanda at 16%, which is where *k13* R561H occurs (*30*). It was postulated that resistance would emerge later in Africa than in Asia because of the higher transmission rate and higher level of immunity (*32*, *33*).

The reduced endocytosis in ART resistant parasites deprives the parasite of amino acids (*34*) and there is clear evidence for a fitness cost of ART resistance (*9*, *35*). Usually considerably less than 50% of parasites from resistant patient isolates survive in the RSA (*8*) and the delayed clearance rather than full loss of susceptibility in patients could indicate that it is possible for parasites with higher levels of ART resistance to arise. However, it is at present unclear if and to what extent the fitness cost impedes higher levels of resistance. It is also unclear if combinations of mutations could lead to hyper-resistant parasites.

Here we test 135 mutations in proteins from the K13-compartment including 4 mutations in K13 itself and find that none of them confers significant levels of resistance over GFP-tagged wild type (WT) K13. However, some of the *k13* mutations resulted in a significantly lower susceptibility to ART than the 3D7 background. Together with known high resistance mutations this provided us with 3D7-based laboratory lines with different resistance levels. A comparison of parasites with different resistance levels showed a correlation between resistance, fitness cost and K13 protein abundance of a given variant, suggesting a constraint of resistance levels by the fitness cost. This was confirmed using parasites artificially selected for higher resistance. We also assess the threat of hyper-resistance, the impact on fitness cost on this and the potential impact of *k13* double mutations and combinations with resistance mutations outside *k13*.

## Results

### Tested mutations in K13-compartment proteins do not cause *in vitro* ART resistance

We aimed to assess whether further mutations in addition to the already identified resistance mutations could cause reduced ART susceptibility. As not only *k13*, but also genes of K13 compartment proteins such as *ubp1* can cause reduced susceptibility, we analyzed non-synonymous mutations in *k13* and genes of other K13 compartment proteins from African patient samples available from WWARN (*36*), the Ghanaian Fever Without Source study (*37*) and MalariaGen (*38*). These included mutations in the genes encoding K13, KIC1, KIC2, KIC4, KIC5, KIC7, KIC9, UBP1, EPS15and MyoF (previously annotated as MyoC) (list of mutations in Table S1). In addition we selected one untested mutation in AP2α, which was reported to be present in *in vitro* ART-selected parasites that displayed a K13-independent reduced ART-susceptibility (*39*). Due to the large number of mutations in the K13 compartment proteins (a total of 131 mutations in ten genes) we created parasites carrying multiple mutations (multi-mutants) by introducing a recodonized sequence of the target gene that encodes several mutations in the same gene into 3D7 parasites using selection-linked integration (SLI, (*40*)) (Table S1). In total 125 mutations were included in 8 multi-mutants and 6 mutations were tested individually (Figure 1A,B). RSAs (using dihydroartemisinin (DHA)) with multi-mutants of compartment members of K13 showed no change in susceptibility to ART, suggesting that none of the mutations in the genes encoding KIC1, KIC2, KIC4, KIC5, KIC7, KIC9, UBP1, EPS15 and MyoF resulted in ART resistance (Figure 1A,B). One exception was AP2α H817P which significantly increased RSA survival, although this was not above 1% which is considered the cut-off for clinically relevant resistance (*7*). Yet this increase might explain the occurrence of this mutation in parasites that had been selected for ART-resistance through ART exposure *in vitro* but that lacked K13 mutations (*39*).

**Figure 1.**
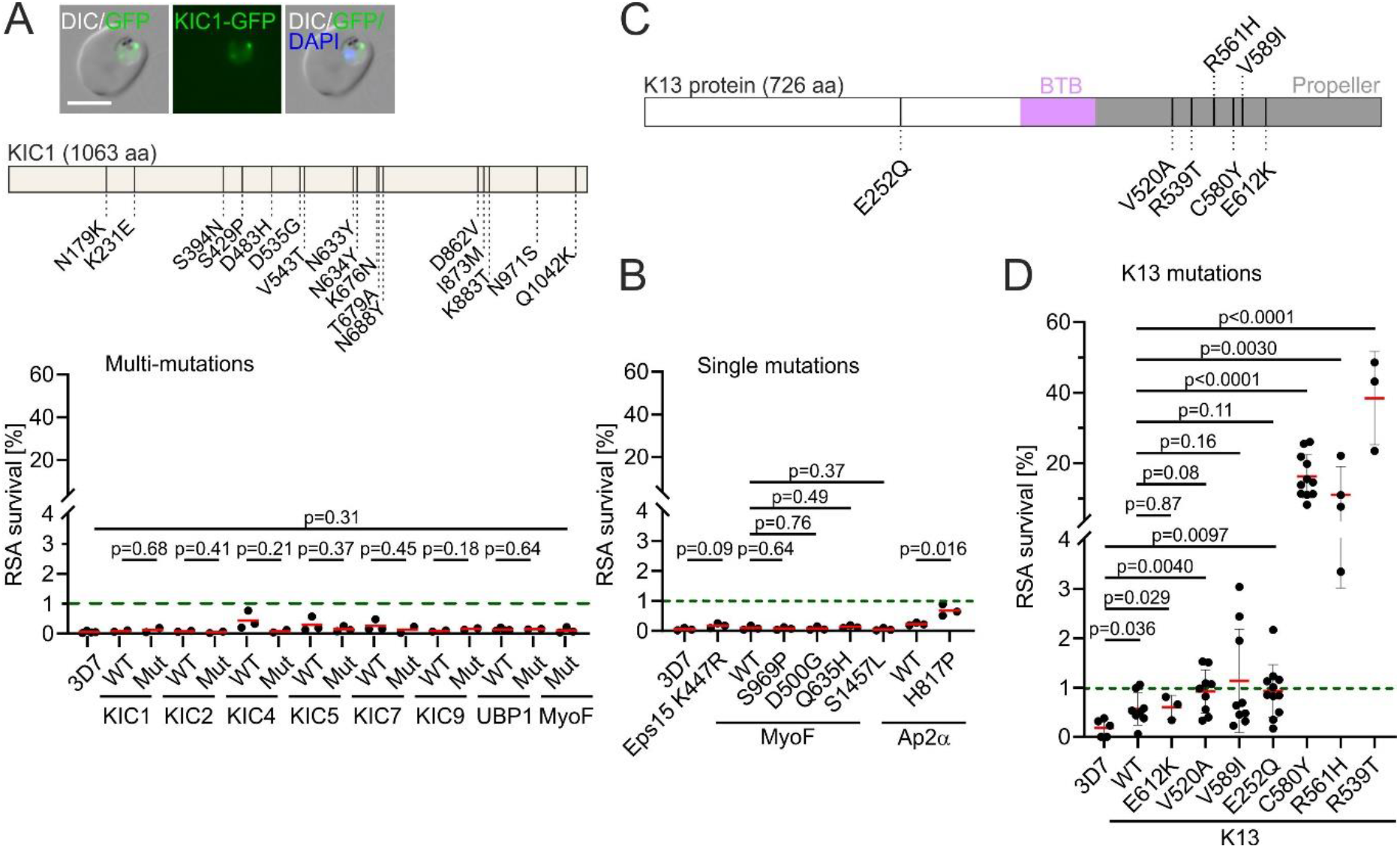
135 tested mutations in K13-compartment proteins do not cause ART resistance. **(A)** Mutations introduced in the multi-mutant of KIC1 are exemplarily shown in a schematic and fluorescence images of the resulting parasites harboring multi-mutant-KIC1 are shown. Scale bar 5 μm. RSA survival for all multi-mutants is displayed (% survival compared to control without DHA 66 h after 6 h DHA treatment in RSA). MyoF multi-mutant was compared with 3D7 as matching WT was not available. **(B)** RSA survival for single mutations in K13 compartment proteins. Eps15 K447R was compared with 3D7 as matching WT was not available. **(C)** Position of *k13* mutations as well as BTB domain and propeller domain are shown on K13. **(D)** RSA of different *k13* mutant cell lines. Error bars show standard deviations. WT, wild type; red bars indicate the mean parasite survival rate of the respective cell line, each dot represents an independent experiment, green dashed line represents the 1% cut-off value defining clinically-relevant ART resistance. Position of *k13* mutations is shown on K13. Error bars show standard deviations. P values derive from two-tailed unpaired t-tests.

For *k13* itself we investigated the mutations V520A, V589I and E612K that have been found with low to moderate prevalence (0.8%-5.0%) in different malaria endemic regions in Africa and were not previously tested for ART resistance *in vitro* (Figure 1C). All mutations originated from studies where the samples had been taken after ACT started to be used in the respective country, with exception of one study in Kenya in 2002 (Dataset S1). We noticed that *k13* V520A and V589I occurred in regions with above average and average malaria incidence, respectively, while *k13* E612K, C580Y, R539T and *k13* R561H, which was recently detected in Rwanda, occurred in regions with below average malaria incidences (Figure S1, Dataset S2). E252Q occurred in Thailand and Myanmar between 2005 and 2013 where it was previously associated with slow parasite clearance (*36*) but not tested *in vitro*. Although only detected in Asia, we included this mutation because it might have been missed in Africa as typically only the propeller domain (amino acid residues 444–721) is sequenced when screening for *k13* mutations. The respective mutations were introduced into the *k13* locus of *P. falciparum* 3D7 parasites together with fluorescent protein GFP to result in a GFP-K13 fusion as done previously (*10*). The susceptibility of these parasites to ART was tested by RSA. Introduction of the GFP to the *k13* locus already led to small but significant reduction in ART-susceptibility (0.2±0.2% vs 0.6% ±0.3%) (K13-WT, Figure 1D), suggesting that addition of the GFP to the protein’s N-terminus resulted in a very mild reduction in K13 activity. All three mutations caused a small further decrease in ART-susceptibility although this was not significantly different over *k13* WT but for *k13* V520A and E252Q was significant over the 3D7 parent (Figure 1D). We conclude that these mutations have likely no relevance for ART-resistance in endemic settings but provide us with parasites with a mildly reduced ART-susceptibility compared with 3D7.

For comparison, we also tested *k13* C580Y, R561H and R539T, which were previously shown to be resistance mutations. Parasites with C580Y (16.3% ±6.1%), R561H (11.1% ±8.0%) and R539T (38.4% ±13.2%) *k13* showed higher RSA survival (Figure 1C), similar to what was previously observed for C580Y and R539T in 3D7 (*8*).

### Double mutations in *k13* do not significantly increase ART resistance

To anticipate how *k13* resistance phenotypes could further develop in endemic areas, we investigated whether combination of the *k13* C580Y mutation (which is highly prevalent in SE Asia) with the V520A mutation might potentiate resistance e.g. through a synergetic effect. Further we investigated whether the two known resistance mutations *k13* C580Y and R539T could combine to result in hyper-resistance. To obtain the corresponding cell lines, we again modified the endogenous *kelch13* locus of 3D7 parasites using SLI so that the double-mutated K13 was fused to GFP. The K13^V520A+C580Y^ parasites did not display an increased resistance level in RSAs (11.5% survival) compared with parasites with only the C580Y mutation (16.3% survival) (Figure 2A), indicating that there is no synergistic effect. Similarly, K13^R539T+C580Y^ parasites showed no significant change in RSA survival (33.4% survival) compared with parasites harboring R539T alone (38.4% survival) but significantly more survival than the C580Y alone, suggesting no additive or synergistic effect. In conclusion, —at least with the tested mutations — there is no indication for additive or synergistic effects.

**Figure 2.**
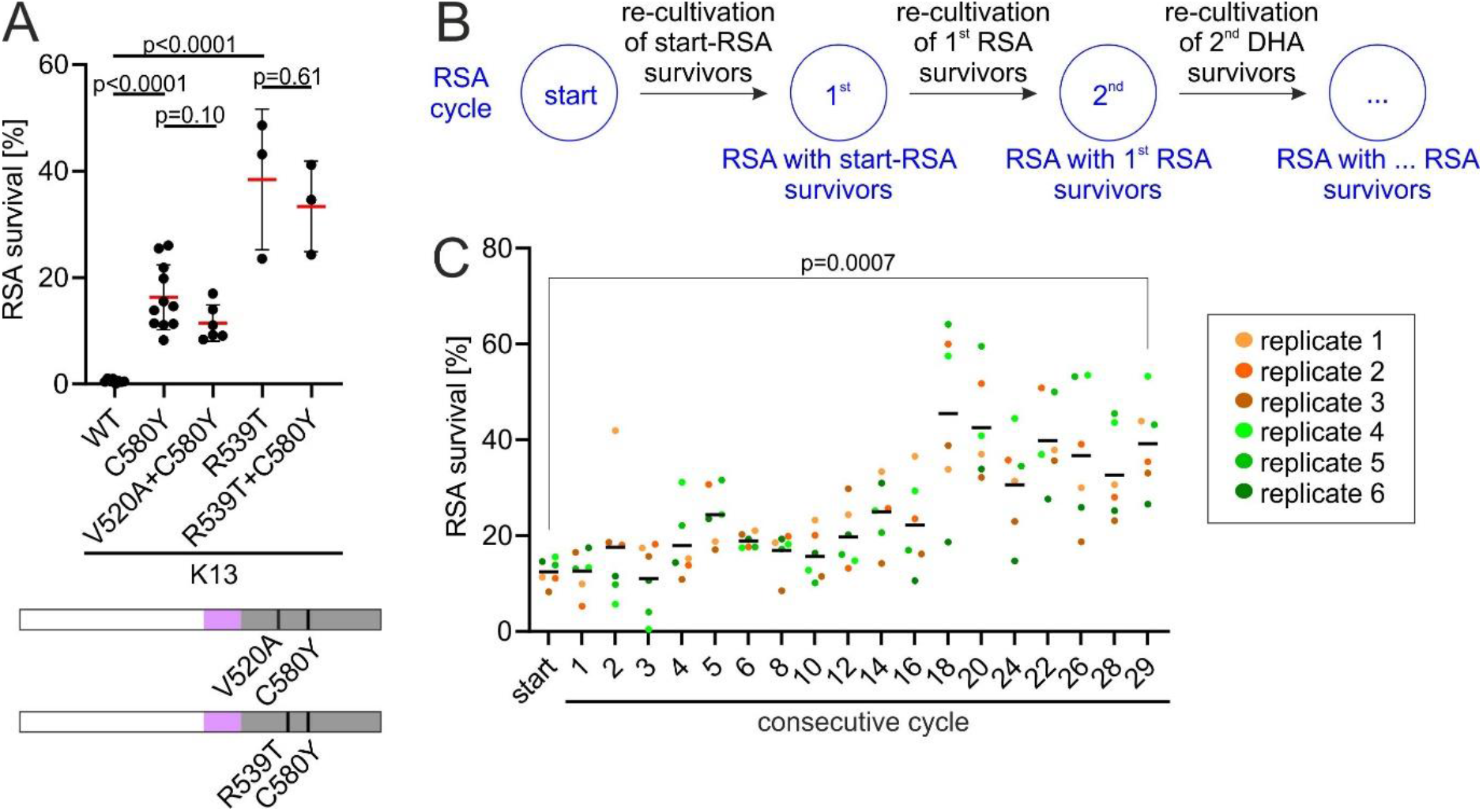
DHA-selection but not combination of two mutations increases resistance. **(A)** RSA survival of the different *k13* mutant cell lines shown. WT, wild type; each point represents an independent RSA; red bars indicate the mean parasite survival rate of the respective cell line. Position of *k13* mutations are shown on K13. Domains are colored as in Figure 1. Red bars show mean; error bars show standard deviations. P values derive from two-tailed unpaired t-tests. **(B)** Scheme of experimental procedure of consecutive RSA cycles performed with DHA survivors of the respective prior cycle. **(C)** Parasite survival rate of K13^C580Y^ (% survival compared to control without DHA) 66 h after 6 h DHA treatment in RSA. Six experiments per cycle were performed, consisting of six replicates (see color code). Replicates 1, 2, 3 (orange) and 4, 5, 6 (green) were each started on the same days. Black bars show mean. P value is indicated, two-tailed paired t-test.

### Selection by consecutive RSAs significantly increases ART resistance in *k13* C580Y parasites

In a further attempt to anticipate increased resistance, we tested whether parasites with a common resistance mutation can be selected for increased resistance by performing consecutive RSAs. For this, an RSA was carried out with K13^C580Y^ parasites and surviving parasites were subjected to a consecutive RSA as soon as they reached sufficient parasitemia following the previous RSA (Figure 2B). After 29 iterations of RSAs, we obtained the parasite line K13^C580Y^-29^th^. These parasites displayed significantly increased survival in RSA (39.2%) compared with the original K13^C580Y^ parasites at the start of this experiment (12.4%) (Figure 2C), showing that repeated ART-treatment of *k13* C580Y-harbouring parasites can increase ART resistance. Sequencing of *k13* in these parasites showed that this was not due to further mutations in *k13* in addition to the C580Y already present at the beginning of the experiment (Figure S2).

### Cellular levels of K13 and KIC7 proteins determine ART-resistance

Previously, we and others showed that the level of K13 in the K13^C580Y^ and K13^R539T^ parasites is lower than in K13^wt^ parasites and that reducing K13 abundance resulted in resistance (*10*, *16*, *18*). We assessed the cellular K13 levels in the parasites for all *k13* mutations in this study, to find whether a change in K13 level generally accompanies resistance mutations. We previously used either Western blots or direct microscopy-based measurement of GFP fluorescence in the cells to determine K13 levels (*10*). Here we used direct microscopy-based measurement of K13 levels because in comparison to Western blots, which are only semi-quantitative, this procedure is not susceptible to influence of cell lysis and the multiple steps required for blotting and detection. These experiments showed no significant reduction in the amount of K13 with the V520A (94% of WT) and the V589I (91% of WT) mutations (Figure 3A). The E252Q mutation showed 105% of the amount (non-significant) of WT (Figure 3A). The resistance mutations *k13* C580Y, *k13* R561H and *k13* R539T had 52%, 51% and 51% of the K13 amount of WT, respectively (Figure 3A). As expected from their RSA resistance behavior, parasites with the *k13* V520A C580Y double mutation harbored similar levels of K13 (48% of WT) as parasites with the *k13* C580Y alone, and parasites with *k13* R539T and C580Y (57% of WT) had similar levels as both *k13* C580Y and *k13* R539T individually. The K13 levels in K13^C580Y^-29^th^ parasites showed a further reduction (32% of WT) when compared with K13^C580Y^ parasites, even though this reduction was not significant when comparing means (Figure 3A), only when comparing all data points (Figure S3). We plotted the cellular K13 levels against RSA survival (*in vitro* resistance). Cellular K13 levels correlated with *in vitro* resistance (Pearson’s r=-0.81, p=0.005) (Figure 3B). We note that this was largely because the mutations with low K13 levels clustered into a group conferring significant *in vitro* resistance over GFP-K13-WT and the other mutations into a group with high K13 levels (Figure 3B).

**Figure 3.**
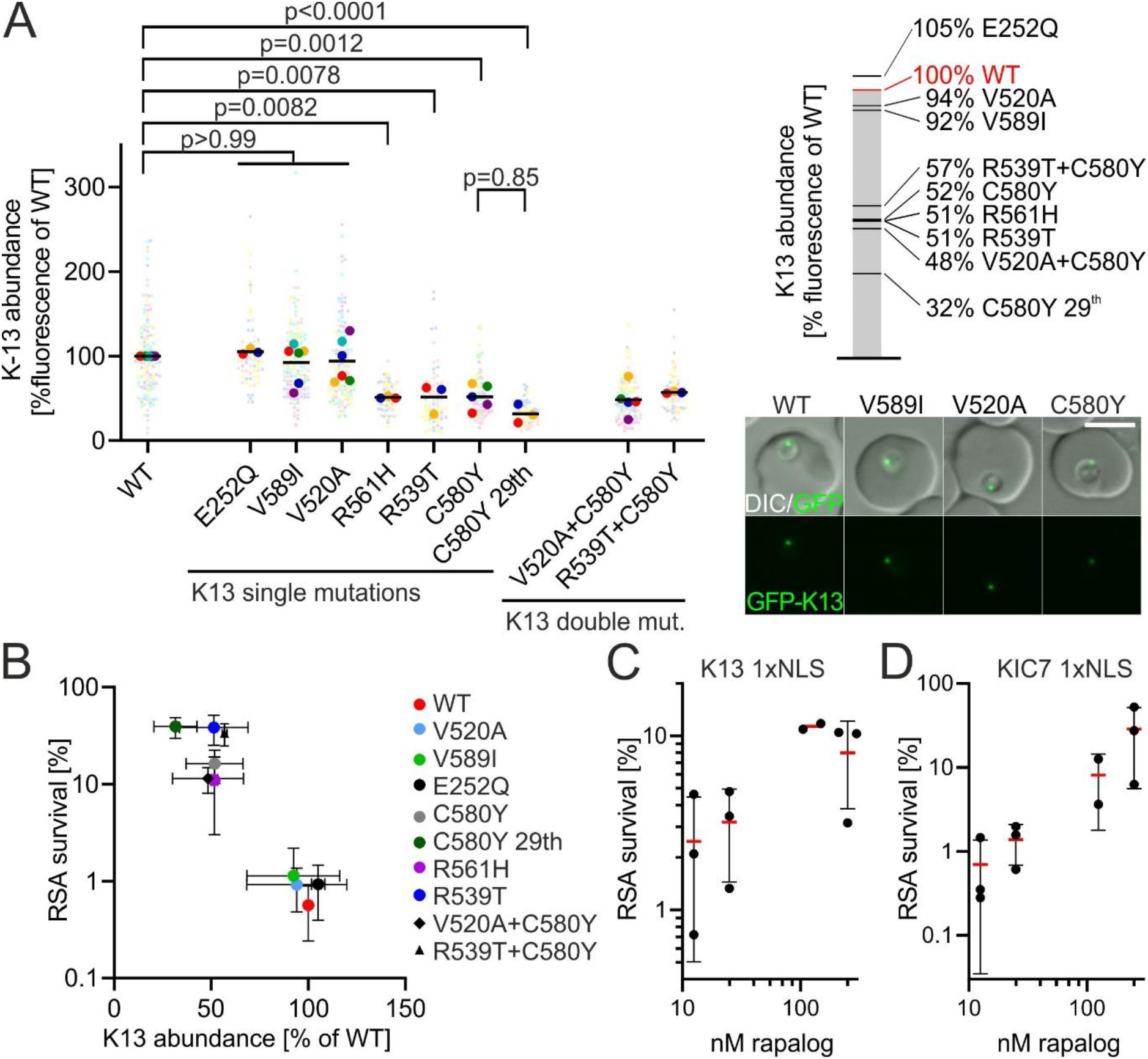
Resistance is inversely correlated with K13- and KIC7-abundance. **(A)** K13-abundance measured by the GFP fluorescence intensity of the single focus observed in ring stage parasites with the indicated K13 expressed from the endogenous locus and fused to GFP, normalized to the fluorescence in parasites with the identically modified endogenous locus but with a WT GFP-K13. Each small dot represents the measured value from one focus in one parasite. Large dots of the same color as small dots represent mean of the respective measured values and each large dot represents one biological replicate. Each biological replicate (large dot) consists of 20 individual measurements (small dots). P-values derive from comparing means (large dots) by one-way ANOVA. Example fluorescence microscopy images are shown. Scale bar 5 μm. **(B)** Mean of parasite survival in RSA plotted against mean of K13-abundance. Error bars show standard deviations. **(C+D)** RSA survival of (C) K13 1xNLS parasites or (D) KIC7 1xNLS parasites grown in the presence of 12.5 nM, 25 nM, 125 nM or 250 nM rapalog for 3 h before and 6 h during the DHA exposure of the RSA. Red bar, mean; standard deviation indicated; each dot derives from an independent experiment.

To test whether RSA-resistance not just correlates with but is determined by the amount of K13 in its cellular location, we used K13 WT 1xNLS parasites (*10*). In this line K13 protein is removed into the nucleus by a nuclear localization signal (NLS) upon addition of rapalog (termed knock sideways), resulting in conditional reduction of K13 at its cellular location. Previously, 250 nM rapalog was used to achieve a knock sideway of K13 (*40*). Here, we used different concentrations of rapalog to result in different levels of K13 protein remaining at its site of action. ART-resistance, measured by RSA, decreased as the rapalog concentration and hence K13 knock sideways decreased, confirming that RSA-survival is a function of K13 levels in the parasites (Figure 3C). These results confirm observations made on parasites in which the K13 abundance was titrated on mRNA level through the glmS system (*19*). To confirm that this was due to a reduction in endocytosis, we performed a second titration using KIC7, a K13-compartment protein, which was also knocked aside using the same decreased rapalog concentrations as for K13 (Figure 3D). Again, RSA survival was dependent on the level of knock sideways of KIC7 (i.e. amount of KIC7 at the K13 compartment), indicating that the activity of the endocytic process determines resistance and that this occurs through the available amount of some of the proteins involved in this process.

### Fitness cost correlates with ART-resistance

Several studies modelling fitness and within-host competition have highlighted the importance of parasite fitness and propose that resistant parasites with reduced fitness are less likely to establish themselves in high transmission areas like most of Africa (*33*, *41*, *42*). Some K13 resistance mutations, including C580Y and R561H, have been shown to have a fitness costs (*22*, *43*). To measure the impact of *k13* mutations on fitness of our parasites with isogenic backgrounds in this work, we first monitored *in vitro* growth over 96 h. This confirmed the fitness cost of *k13* C580Y and suggested that generally, fitness cost might correlate with resistance but the variation between experiments was too high for reliable conclusions (Figure 4A,B).

**Figure 4.**
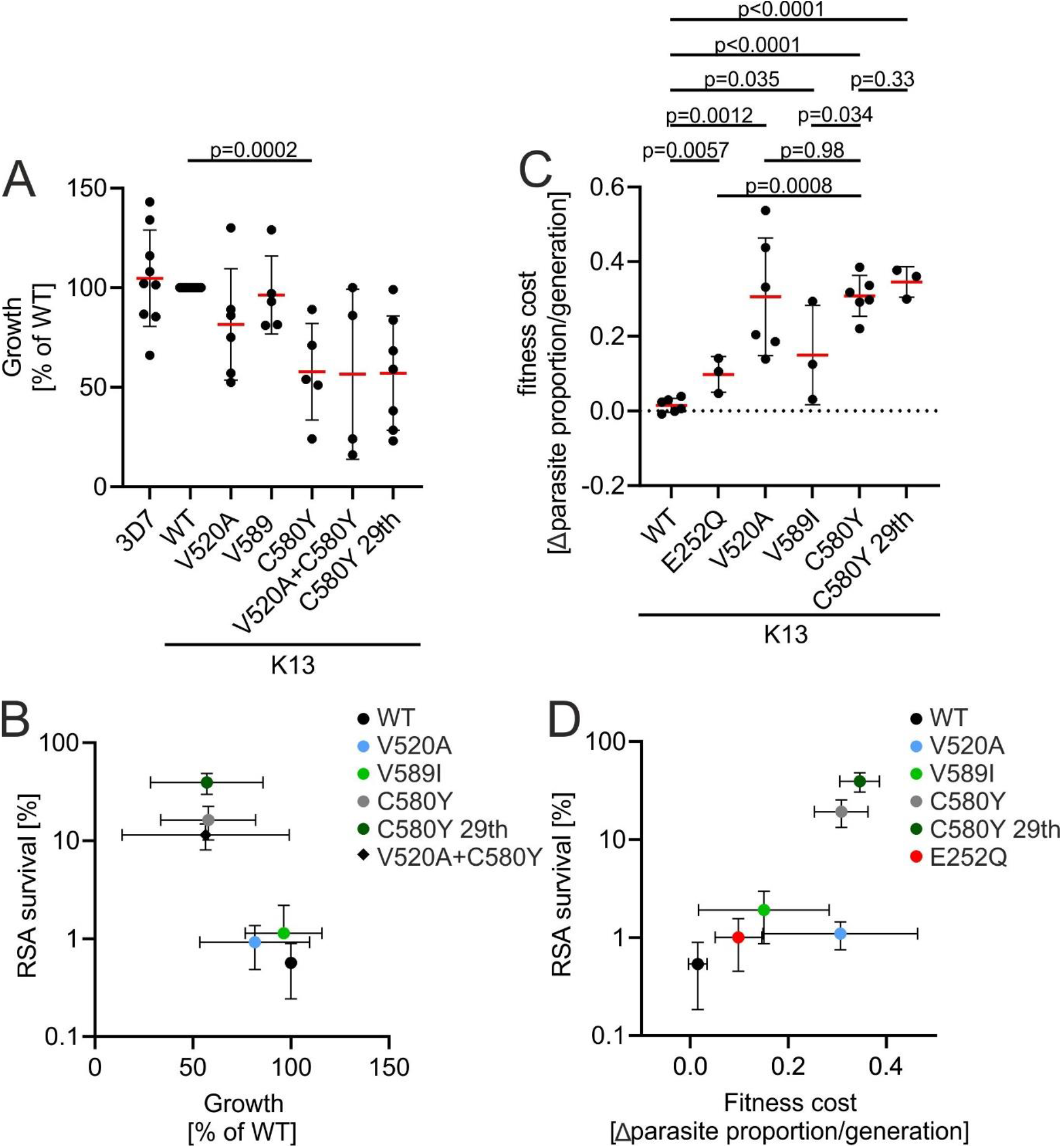
Fitness cost correlates with resistance: **(A)** Parasite growth, calculated as fold-change of parasitemia after 96 h compared to parasitemia at 0 h, normalized to WT growth for different cell lines with indicated k13 mutations. Error bars show standard deviation. P values derive from two-tailed unpaired t-tests. **(B)** Mean of growth of strains harboring different *k13* mutations plotted against mean of parasite survival in RSA. Error bars show standard deviation. **(C)** Fitness cost of parasites harboring different *k13* mutations grown in competition with 3D7 parasites given as loss of parasite proportion/generation. Error bars show standard deviation. P values derive from two-tailed unpaired t-tests. **(D)** Mean of fitness cost plotted against mean of parasite survival in RSA. All error bars show standard deviations.

For better fitness cost measurements, parasite proportions were determined in a mixed culture of a mutant and the 3D7 parent parasite line, as previously described (*22*, *44*). Parasites were mixed 1:1 with the 3D7 parasites and the proportion of mutated (fluorescent) parasites was tracked until it made up less than 5% of the parasite population (Figure 4C, S3). The parasites with K13 mutations (V520A, V589I, E252Q, C580Y and C580Y-29th) declined in proportion compared to 3D7 in all 6 experiments whereas GFP-K13 WT did not or only to a small extent (Figure S3). Similar to what was observed when parasites growth was tracked in individual cultures, there was a trend that the RSA-survival caused by a mutation correlated with the fitness cost inflicted by the mutation (Figure 4D). However, this trend was not significant (Pearson’s r=0.69, p=0.13) because *k13* V520A behaved as an outlier in half of the experimental repeats, in which its growth slowed severely after the start of the experiment (Figure S4), while in the other half it grew in line with the trend of the other mutations (Pearson’s r=0.90, p=0.04 when V520A was entirely excluded) (Figure 4C,D). Together with the findings on K13 abundance (Figure 3), it is therefore likely that both fitness cost and ART-resistance are caused by the reduced amount of K13 in the cell as a result of the destabilization of the K13 protein through the given mutation.

### *ubp1* R3138H has a disproportional fitness cost that is further aggravated in K13^C580Y^ parasites

After assessing the effect of the *k13* double mutations V520A+C580Y and R539T+C580Y, we wondered whether there are combinations of mutations or gene disruptions beyond *k13* that harbour the risk of very high resistance levels. We tested this by combining *k13* C580Y with *ubp1* R3138H and the resulting K13^C580Y^+UBP1^R3138H^ parasites did not display a significantly increased resistance level (17.3% mean survival) compared with parasites with the C580Y mutation alone (Figure 5A). We also attempted to disrupt both *kic4* and *kic5*, two genes we were previously able to disrupt individually (*10*), or combine their disruption with UBP1^R3138H^ but were unsuccessful despite six attempts per combination, which may indicate that the fitness cost of combining these prevented us from obtaining these parasite lines.

**Figure 5.**
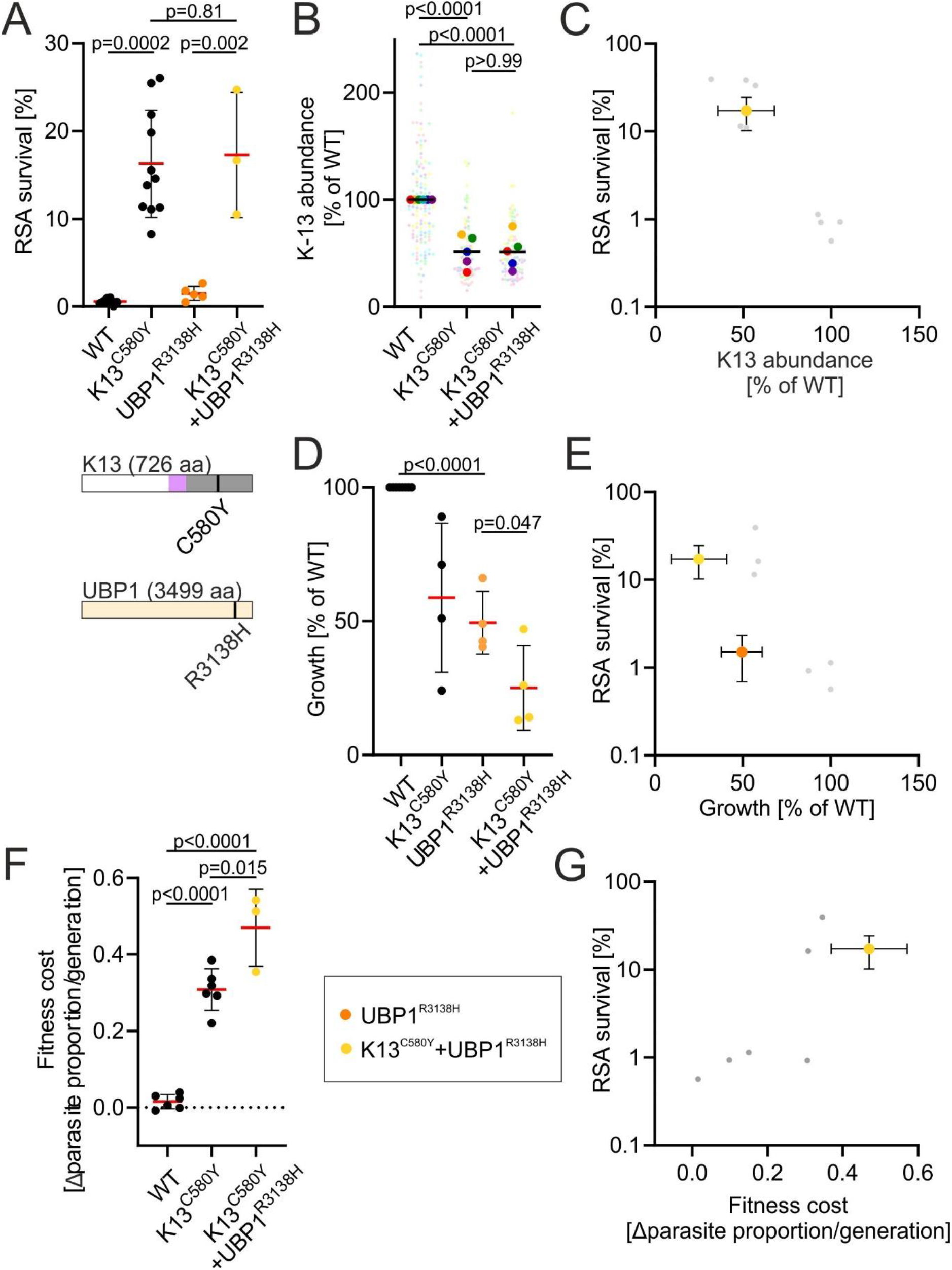
Double mutation of *k13* C580Y and *ubp1* R3138H result in a high fitness cost. **(A)** RSA of K13^C580Y^ and UBP1^R3138H^ single and double mutant cell lines. Red bars indicate the mean parasite survival rate of the respective cell line. Data for UBP1^R3138H^ was previously published in (*10*). Position of *k13* and *ubp1* mutation is depicted in K13 and UBP1. K13 domains are colored as in Figure 1. Error bars show standard deviation. P values derive from two-tailed unpaired t-tests. **(B)** K13-abundance measured GFP fluorescence intensity of the single focus observed in ring stage parasites of the indicated K13 expressed from the endogenous locus and fused to GFP, normalized to the fluorescence in parasites with the identically modified endogenous locus but with a WT GFP-K13. Small dots represent measured values for individual parasites. Large dots of the same color as small dots represent mean of the respective measured values and each large dot represents one biological replicate. Dots of the same color were obtained as part of the same experimental repeat. P values derive from two-tailed unpaired t-tests. **(C)** K13 abundance plotted against parasite survival in RSA for K13C580Y+UBP1R3138H parasites (yellow). Grey dots show data of k13 mutations from Figure 3B for comparison. Error bars show standard deviation. **(D)** Growth, calculated as fold-change of parasitemia after 96 h normalized compared to parasitemia at 0 h for different cell lines with indicated *k13* or *ubp1* mutations. Error bars show standard deviation. P values derive from two-tailed unpaired t-tests. **(E)** Growth of strains harboring *ubp1* R3138H with (yellow) and without (orange) *k13* C580Y plotted against parasite survival in RSA. Grey dots show data of *k13* mutations from Figure 4B for comparison. Error bars show standard deviation. **(F)** Fitness cost for different cell lines with indicated *k13* or *ubp1* mutations grown in mixed cultures with 3D7. Error bars show standard deviation. P values derive from two-tailed unpaired t-tests. **(G)** Fitness cost K13^C580Y^+UBP1^R3138H^ parasites (yellow) plotted against parasite survival in RSA. Grey dots show data of *k13* mutations from Figure 4D for comparison. All error bars show standard deviations.

We further characterized the K13^C580Y^+UBP1^R3138H^ parasites and found that they displayed a similar K13 abundance to resistance level ratio (Figure 5B,C) as the *k13* mutation (Figure 3B). The growth of K13^C580Y^+UBP1^R3138H^ parasites was slower than that of K13^C580Y^ parasites and UBP1^R3138H^ parasites, which each confer a fitness cost, both when measured in individual cultures and direct competition assays (Figure 5D-G). The ratio of fitness cost to resistance level was higher than for *k13* mutations alone (Figure 5E,G). To determine whether just of the combination of the two mutations led to such a reduced growth rate or whether it was due to *ubp1* R3138H by itself, we determined the growth rate of UBP1^R3138H^ parasites. Interestingly, already the *ubp1* R3138H mutation alone caused a disproportionally low growth compared with the level of ART resistance it affords when compared to all tested *k13* mutations (Figure 5E). This is congruent with our previous finding that in contrast to K13 which is only required in rings, UBP1 is also important for hemoglobin endocytosis in trophozoites (*10*) and could explain why we were not able to combine this mutation with *kic4* and *kic5* TGDs and why this mutation has rarely been observed in patient samples. Overall, due to its disproportionally high fitness cost and lacking synergism with *k13* mutations, *ubp1* R3138H and possible mutations in other K13 compartment proteins important for endocytosis in trophozoites, are therefore unlikely to form combinations with *k13* mutations in endemic countries.

## Discussion

In this study we tested 131 mutations in ten genes and found that none of the tested mutations in the genes encoding KIC1, KIC2, KIC4, KIC5, KIC7, KIC9, UBP1, EPS15 and MyoF resulted in significantly decreased susceptibility to ART. We also tested K13 mutations. We found that N-terminally fusing a tag to K13, even without any mutations, very mildly reduced the susceptibility to ART. This effect is negligible compared with the larger effects of common resistance mutations such as C580Y, R561H and R539T and likely does not generally impact experiments with these parasites but needs to considered when assessing mutations with small effect sizes. Thus, we found that the four previously unstudied *k13* mutations did not cause significant resistance in the RSA compared with K13-WT. The small non-significant decrease in susceptibility that was observed for *k13* V520A, V589I and E252Q likely has no clinical impact but we can’t exclude that these variants have some small advantage in the population that is difficult to assess. However, all of these mutations that were tested in competition assays were consistently outcompeted by 3D7 whereas GFP-K13 WT was not, indicating that they have a small disadvantage over WT. Despite likely not being clinically relevant, parasites with a GFP-fused K13 with or without the V520A, E252Q and E612K were significantly less susceptible to ART in RSA compared to 3D7 and together with the high resistance parasites offered us a tool to explore the properties of parasites with different levels of *in vitro* ART resistance.

We also found that the combination of *k13* C580Y with *k13* V520A, or of two resistance mutations with each other, like *k13* C580Y and *k13* R539T, and *k13* C580Y with *ubp1* R3138H did not increase ART resistance. These are the first combinations of C580Y with other naturally occuring mutations generated in isogenic backgrounds. A similar non-additive result was obtained in a recent study for *coronin* R100K and E107V, two *in vitro* selected mutations, combined with *k13* C580Y (*14*).

Previous work has shown that the genetic background can influence the resistance level of parasites with identical *k13* resistance alleles (*9*, *22*), indicating that mutations outside the K13 compartment exist that modulate the ART susceptibility of parasites with *k13* resistance mutations and remain to be determined. Repeated ART-treatment in consecutive RSAs of *k13* C580Y-harbouring parasites increased ART resistance, but this was not due to additional *k13* mutations apart from C580Y. Parasites from such selections might thus be a promising tool to study factors outside of *k13* that influence resistance levels in parasites with a resistance-conferring allel of *k13*.

We found that the amount of K13 and KIC7 at the K13 compartment in the cell determine the level of *in vitro* resistance, confirming that the endocytic process is important for ART resistance and suggesting that the amount of protein directly influences the rate of endocytosis and level of ART susceptibility. The same effect on resistance was observed in a *k13* mRNA titration recently (*19*) and a role of K13 protein abundance rather than a specific functional change in resistance is also supported by the finding that increasing levels of K13 C580Y reverts resistant parasite back to sensitive (*10*). The finding that KIC7 titration had a similar effect strengthens this conclusion and further supports that resistance is due to altered endocytosis levels. We found that all tested resistance mutations had low K13 levels and all other tested mutations had high K13 levels. The protein levels in the K13 double mutants were comparable to that of the single resistance mutation, indicating that at least in these combinations, multiple mutations do not further destabilize K13. In conclusion, for all tested *k13* resistance mutations, the amount of K13 in the parasite, likely through destabilization of K13, was a factor contributing to or causing the resulting resistance.

We further showed that fitness cost of *k13* mutations generally correlated with resistance. This agrees with a recent study looking at *k13* M579I, C580Y and R561H in several African strains including 3D7 although no such correlation was observed in the Asian strain Dd2 (*22*). It is unclear what caused these differences between African strains and Dd2. Nonetheless, it showed that the genetic background in which the mutations occur can influence the fitness cost and the resistance level (*22*, *35*, *45*). Many differences have been observed in ART resistant parasites compared with sensitive parasites (*17*, *20*, *46–51*) and some of these might correspond to such changes in the background that alter the effect of K13 mutations, although some of these changes likely also are downstream effects of the resistance mechanism (*34*). The data presented here originates from a set of isogenic 3D7-derived parasites which differ from each other only in the studied mutations. While this limits extrapolation of the conclusions to other parasite strains, it presents the advantage of separating effects caused by the studied mutations from effects of the genetic background. Due to its long presence in culture, 3D7 also avoids variations that may arise in more recently culture adapted lines that often show reduced growth levels and may still acquire further changes during continued culture to adapt for better growth in culture.

*K13* E252Q was previously observed to inflict a fitness cost that is smaller than that of *k13* C580Y, however, these experiments were not performed using parasites with isogenic backgrounds (*44*). Our results confirm this finding in an isogenic background. The E252Q mutation caused low but significant *in vitro* resistance over 3D7 but not over WT in our study. It was previously associated with delayed parasite clearance in patients, suggesting that either E252Q can only confer resistance in a specific parasite background or that it was not the cause of the delayed clearance. Regarding K13^C580Y^-29^th^, it should also be noted that the long duration of the competition experiments may permit the K13^C580Y^-29^th^ to revert back to a state similar to the K13^C580Y^ parasites, as the continuous RSA selection pressure had to be lifted in the competition assay.

We observed a disproportionally high fitness cost in parasites with the *ubp1* mutation conferring ART resistance. The high fitness cost is likely due to its importance for hemoglobin endocytosis in trophozoites. This is in agreement with the finding that lowering endocytosis by conditional inactivation or disruption of K13 compartment proteins impairs parasite growth (*10*, *52*) and that parasites with a *k13* ART resistance mutation are hypersensitive to low amino acids (*34*). The small fitness cost of *k13* mutations compared to other K13-compartment proteins is likely due to its role in endocytosis exclusively in rings (*10*). Our findings therefore highlight the unique property of K13 which stands in contrast to resistance-conferring changes in KICs that also affect endocytosis in later stage parasites, thus incurring a higher fitness cost. This is a likely reason why *k13* is the predominant gene mutated to cause delayed clearance in patients. Due to their high fitness cost, changes outside *k13* are less likely to arise and the resistance level, as observed with the few found in the field (*15*, *53*, *54*), is low. The high fitness cost of mutations of proteins involved in endocytosis outside *k13* also indicates that combinations of them with *k13* resistance mutations are unlikely to lead to hyper-resistant parasites in the field. We also did not detect any indication for additive or synergistic effects on *in vitro* ART resistance when the *ubp1* resistance mutation and *k13* C580Y were both present in the parasite or if two *k13* mutations were combined.

Our findings also indicate that there seems to be no danger of hyper resistant parasites due to a combination of the two most common resistance mutations *k13* C580Y and *k13* R539T, although we cannot exclude that other combinations of mutations lead to increased resistance. Based on the one tested combination it would also indicate that mutations with minimal increase in ART resistance such as V520A are not a stepping stone for potentiated resistance when combined with current resistance mutations. The only means by which we were able to increase *in vitro* ART resistance was a constant RSA selection regime. As in our isogenic parasites, higher resistance was accompanied by higher fitness costs, it is possible that fitness cost limits the level of ART resistance parasites can reach.

## Materials and Methods

### Plasmid construction

All primers used in this study are listed in Dataset S3. The mutations tested in *k13* and *k13* compartment proteins were chosen to include only mutations that were present in a maximum of 5% of samples taken in the sampling study. For the multi-mutants the pSLI-TGD plasmid (*40*) was used. The homology region was directly linked to a synthesized functional codon-changed version of the corresponding candidate containing all selected mutations (Genscript). The fragments of the homology region and the mutated recodonized sequence of all candidates were cloned by Gibson assembly into the pSLI-TGD vector via NotI/MluI, resulting in the vectors pSLI-KIC1mutpool, pSLI-KIC2mutpool, pSLI-KIC4mutpool, pSLI-KIC5mutpool, pSLI-KIC7mutpool, pSLI-KIC9mutpool, pSLI-UBP1mutpool and pSLI-MyosinFmutpool. Sequencing was performed to confirm absence of undesired mutations.

The N-terminal SLI plasmid of K13, pSLI-N-GFP-2xFKBP-K13-loxP (*40*) was modified for the Kelch13 mutant parasites as follows: the mutation was obtained by amplifying the codon-changed synthesized version of *k13* using primers encoding the mutation. The resulting two fragments were cloned into the pSLI-N-GFP-2xFKBP-K13-loxP via AvrII/StuI.

To generate K13^V520A+C580Y^ and K13^R539T+C580Y^, pSLI-N-GFP-2xFKBP-K13_C580Y-loxP (*40*) was modified by amplifying *k13* from this plasmid using primers to introduce the mutation and ligating the resulting fragments into the same vector at the AvrII/XhoI sites.

The *k13* C580Y mutation was introduced into UBP1^R3138H^ parasites using a SLI2a plasmid based on the pSLI-N plasmid (*40*) but containing the BSD resistance gene instead of the hDHFR gene (*55*). The hDHFR was excised using BamHI and HindIII. KIC4-TGD and KIC5-TGD plasmids were created by inserting the respective homology regions (*10*) into SLI2a plasmids based on the pSLI-TGD plasmid (*55*).

### Parasite culturing and transfection

*P. falciparum* 3D7 parasites (*56*) were cultivated at 37°C in 0+ erythrocytes in RPMI complete medium with 0.5% Albumax (Life Technologies) with 5% hematocrit and transfected as previously described (*57*, *58*). All parasite lines used in this study are listed (Table S2). Transgenic parasites were selected using 4 nM WR99210 (Jacobus Pharmaceuticals) or 2.5 μg-ml BSD (Invitrogen). For selection-linked integration, parasites were selected as previously described using 0.9 μM DSM1 (BEI resources) or 400 μg/ml G418 (Merck) (*40*). Correct integration was confirmed by PCR as described (*40*) (Figure S5). Parasite line-specific integration check primers are listed in Dataset S3.

### *In-vitro* ring-stage survival assay^0-3h^ (RSA) and consecutive RSAs

All RSAs were performed according to the standard procedure described previously (*59*). 0-3 h old rings were treated with 700 nM DHA for 6 h and cultivated for another 66 h at 37°C. Giemsa smears were taken and parasite survival rate determined by comparing the parasitemia of viable parasites after DHA against the parasitemia of the untreated control. Parasites were defined as resistant when mean survival rate exceeded the cut-off value of 1% (*59*).

For the consecutive RSA, K13^C580Y^ parasites (*10*) were used. 66 h after the DHA pulse, Giemsa smears were taken and the surviving parasites of the DHA treated sample were re-cultivated in a new Petri dish. After sufficient parasitemia was reached, the parasites originating from the RSA survivors were subjected to a new RSA. This procedure was continuously repeated for 30 rounds. The *k13* gene of the resulting parasites K13^C580Y^-29^th^ was sequenced and showed no changed compared to the starting cell line which harbored a recodonized *k13* with the C580Y mutation (*10*).

### Fluorescence microscopy

Microscopy was performed as described earlier (*60*). A Zeiss Axio Imager M1 or M2 provided with a Hamamatsu Orca C4742-95 camera was used for imaging. Zeiss Plan-apochromat 63x or 100x oil immersion objectives with 1.4 numerical aperture were used. Images were edited using Corel Photo Paint X8 and brightness and intensity were adjusted. Images that were used for quantification were not adjusted for brightness and intensity.

### Measurement of protein amount by fluorescence intensity

GFP-K13 parasites were synchronized two times using 5% sorbitol at intervals of two days. After the second sorbitol synchronization, the cell lines were cultivated at 37°C for 2 more hours and then GFP signal of the ring-stage parasites was detected by fluorescence microscopy using the 63x oil immersion objective. GFP-K13 WT parasites were always imaged alongside parasites carrying mutations and were used to normalize the signal of mutation-harboring parasites. Parasites were selected based on DIC and then exposed for 200 ms to image green fluorescence. Total intensity of the GFP signal in foci was measured and background signal subtracted using ImageJ (ImageJ2 2018, (*61*)).

### Growth assessment

The parasitemia of a mixed-stage parasite culture was measured by flow cytometry (*40*) and based on this the parasitemia was adjusted to 0.05 to 0.1% parasitemia in 2.5% hematocrit. The parasitemia was then measured again by flow cytometry to determine the start parasitemia. These parasites were cultivated for 96 h and the medium was changed every 24 h. After 96 h parasitemia was measured again by flow cytometry and divided by the starting parasitemia to obtain the fold change in parasitemia.

### Competition assay

Schizonts were isolated using 60% percoll purification, washed once with medium and cultured for 6 hours to allow invasion of merozoites into new red blood cells. Remaining schizonts were removed by a 10-min incubation in 5% sorbitol solution. The resulting 0-6 h old parasites were cultured at 37°C for 20-24 h after which the parasitemia was measured by flow cytometry (*40*). Based on the determined parasitemia, the mutant K13 cell lines were co-cultivated in a 1:1 ratio with 3D7 control in a 5 mL Petri dish. The proportion of GFP-positive parasites was assessed by fluorescence microscopy until one parasite line reached 95% of parasite proportion.

### Statistical analysis and malaria incidence data

Unpaired or paired two-tailed t-tests were performed as indicated and Pearson’s r calculated using GraphPad Prism 9.0.2. Linear regressions were fit to log transformed data (GraphPad Prism). All error bars shown are standard deviations. Malaria incidence data was downloaded from https://ghdx.healthdata.org/gbd-results-tool.

## Acknowledgments

We thank Jacobus Pharmaceuticals for supplying WR99210. DSM1 (MRA-1161) was received from MR4/BEI Resources, NIAID, NIH. We thank Birgit Förster for sequencing and Ralf Krumpkamp for helpful discussions. This research made use of PlasmoDB.org. HMB acknowledges funding by the Ortrud Mührer Fellowship of the Vereinigung der Freunde des Tropeninstituts Hamburg e.V.. This study was in part funded by the German Center for Infection Research (TTU03.806). OMA was supported by the DELGEME fellowship funded by DELTAS-Africa 107740/Z/15/Z from Wellcome Trust. TS, IH and RS acknowledge funding by the European Research Council (ERC, grant 101021493).

## Author contributions

TS, OMA and JM conceived the project. TS supervised research. SSchm, HMB, RS, IH and DP performed experiments. SSchm and HMB analyzed data and prepared figures. HMB and TS wrote the manuscript draft. All authors approved the manuscript.

## Data and materials availability

All data are available in the main text or the supplementary materials.

