## Supplemental Material for "Impact of different mutations on Kelch13 protein levels, ART resistance and fitness cost in *Plasmodium falciparum* parasites"

**This PDF file includes:**

Figs. S1 to S5
Tables S1 to S2
Dataset S1 to S3

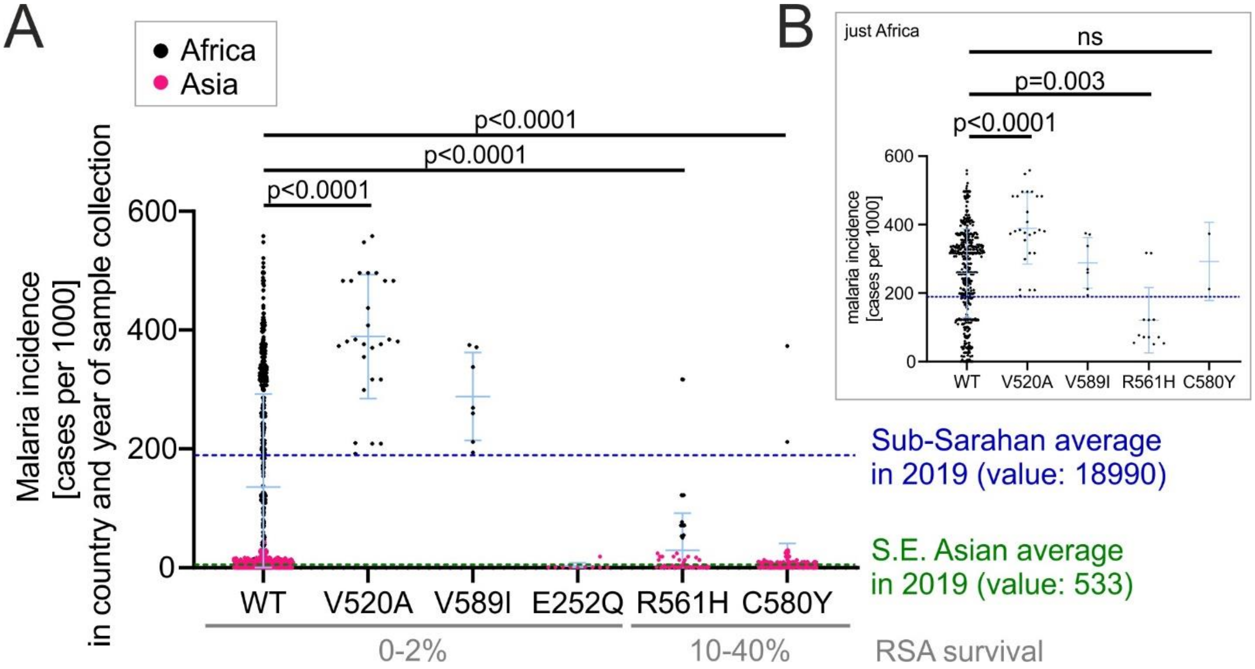


**Fig. S1.**

**Resistance mutations R561H and C580Y are detected at below average malaria incidence. (A)** Malaria incidence at place and time of *k13* mutation detection. Detection of selected k13 mutations in Africa and Asia is shown. Error bars show standard deviation. P-values from unpaired t-tests. Blue dotted line shows average malaria incidence in Sub-Saharan Africa in 2019. Green dotted line shows average malaria incidence in South East Asia in 2019. RSA survival shown as determined in this work (Figure 1B). **(B)** Same data as in A, excluding detection in Asia. P-values from unpaired t-tests of African data only.

```
K13_C580Y_sequence        -----------------------------------ATGGAGGGTGAGAAGGTTAAGACTA    25
sequencing_primer_1       CTGCTGCTGGTGCTGGAGGTGCAGGTAGACCTAGGATGGAGGGTGAGAAGGTTAAGACTA    120


K13_C580Y_sequence        AAGCTAACAGTATAAGTAACTTCAGTATGACTTACGACAGAGAGAGTGGAGGGAATAGTA    85
sequencing_primer_1       AAGCTAACAGTATAAGTAACTTCAGTATGACTTACGACAGAGAGAGTGGAGGGAATAGTA    180

K13_C580Y_sequence        ACTCAGACGACAAGAGTGGTTCAAGTTCAGAAAACGACTCAAACAGTTTTATGAACTTAA    145
sequencing_primer_1       ACTCAGACGACAAGAGTGGTTCAAGTTCAGAAAACGACTCAAACAGTTTTATGAACTTAA    240

K13_C580Y_sequence        CATCTGACAAGAACGAAAAGACAGAGAACAACTCATTTTTACTTAACAACTCATCTTACG    205
sequencing_primer_1       CATCTGACAAGAACGAAAAGACAGAGAACAACTCATTTTTACTTAACAACTCATCTTACG    300

K13_C580Y_sequence        GTAACGTAAAGGACTCTTTATTGGAGTCTATAGACATGTCAGTGTTGGACAGTAATTTCG    265
sequencing_primer_1       GTAACGTAAAGGACTCTTTATTGGAGTCTATAGACATGTCAGTGTTGGACAGTAATTTCG    360

K13_C580Y_sequence        ACTCAAAGAAGGACTTCTTGCCTTCAAACTTGTCTAGGACTTTCAACAACATGAGTAAGG    325
sequencing_primer_1       ACTCAAAGAAGGACTTCTTGCCTTCAAACTTGTCTAGGACTTTCAACAACATGAGTAAGG    420

K13_C580Y_sequence        ACAACATTGGTAACAAGTACTTGAACAAGTTATTGAACAAGAAGAAGGACACAATAACTA    385
sequencing_primer_1       ACAACATTGGTAACAAGTACTTGAACAAGTTATTGAACAAGAAGAAGGACACAATAACTA    480

K13_C580Y_sequence        ACGAGAACAACAACATAAACCACAACAACAACAACAACAACTTAACTGCTAACAACATTA    445
sequencing_primer_1       ACGAGAACAACAACATAAACCACAACAACAACAACAACAACTTAACTGCTAACAACATTA    540

K13_C580Y_sequence        CAAACAACTTAATAAACAACAACATGAACTCACCTTCTATAATGAACACAAATAAGAAGG    505
sequencing_primer_1       CAAACAACTTAATAAACAACAACATGAACTCACCTTCTATAATGAACACAAATAAGAAGG    600

K13_C580Y_sequence        AAAACTTCTTGGACGCTGCTAACTTGATTAACGACGACTCAGGTTTGAATAACTTGAAGA    565
sequencing_primer_1       AAAACTTCTTGGACGCTGCTAACTTGATTAACGACGACTCAGGTTTGAATAACTTGAAGA    660
sequencing_primer_2       ------TCTTGGACGCTGCTAACTTGATTAACGACGACTCAGGTTTGAATAACTTGAAGA    197

K13_C580Y_sequence        AGTTCTCTACAGTTAACAACGTTAACGACACATACGAGAAAAAGATAATAGAGACCGAGT    625
sequencing_primer_1       AGTTCTCTACAGTTAACAACGTTAACGACACATACGAGAAAAAGATAATAGAGACCGAGT    720
sequencing_primer_2       AGTTCTCTACAGTTAACAACGTTAACGACACATACGAGAAAAAGATAATAGAGACCGAGT    257

K13_C580Y_sequence        TGTCAGACGCATCAGACTTCGAGAACATGGTTGGAGACTTGAGGATAACTTTCATAAACT    685
sequencing_primer_1       TGTCAGACGCATCAGACTTCGAGAACATGGTTGGAGACTTGAGGATAACTTTCATAAACT    780
sequencing_primer_2       TGTCAGACGCATCAGACTTCGAGAACATGGTTGGAGACTTGAGGATAACTTTCATAAACT    317

K13_C580Y_sequence        GGTTGAAGAAAACTCAGATGAACTTCATAAGAGAGAAGGACAAGTTGTTCAAGGACAAAA    745
sequencing_primer_1       GGTTGAAGAAAACTCAGATGAACTTCATAAGAGAGAAGGACAAGTTGTTCAAGGACAAAA    840
sequencing_primer_2       GGTTGAAGAAAACTCAGATGAACTTCATAAGAGAGAAGGACAAGTTGTTCAAGGACAAAA    377

K13_C580Y_sequence        AGGAGCTTGAGATGGAGAGGGTTCGTTTATATAAGGAGTTGGAGAATCGTAAGAACATAG    805
sequencing_primer_1       AGGAGCTTGAGATGGAGAGGGTTCGTTTATATAAGGAGTTGGAGAATCGTAAGAACATAG    900
sequencing_primer_2       AGGAGCTTGAGATGGAGAGGGTTCGTTTATATAAGGAGTTGGAGAATCGTAAGAACATAG    437

K13_C580Y_sequence        AGGAGCAAAAGTTGCACGACGAGAGGAAAAAGTTGGACATAGACATTTCAAACGGATACA    865
sequencing_primer_1       AGGAGCAAAAGTTGCACGACGAGAGGAAAAAGTTGGACATAGACATTTCAAACGGATACA    960
sequencing_primer_2       AGGAGCAAAAGTTGCACGACGAGAGGAAAAAGTTGGACATAGACATTTCAAACGGATACA    497

K13_C580Y_sequence        AGCAGATTAAGAAGGAGAAGGAGGAGCACAGAAAGCGTTTCGACGAGGAGAGGTTGAGGT    925
sequencing_primer_1       AGCAGATTAAGAAGGAGAAGGAGGAGCACAGAAAGCGTTTCGACGAGGAGAGGTTGAGGT    1020
sequencing_primer_2       AGCAGATTAAGAAGGAGAAGGAGGAGCACAGAAAGCGTTTCGACGAGGAGAGGTTGAGGT    557

K13_C580Y_sequence        TCTTGCAGGAGATAGACAAGATAAAGTTGGTTTTGTACTTGGAGAAGGAGAAGTACTACC    985
sequencing_primer_1       TCTTGCAGGAGATAGACAAGATAAAGTTGGTTTTGTACTTGGAGAAGGAGAAGTACTACC    1080
sequencing_primer_2       TCTTGCAGGAGATAGACAAGATAAAGTTGGTTTTGTACTTGGAGAAGGAGAAGTACTACC    617

K13_C580Y_sequence        AGGAGTACAAGAACTTCGAAAACGACAAGAAGAAGATAGTAGACGCTAACATAGCAACAG    1045
sequencing_primer_1       AGGAGTACAAGAACTTCGAAAACGACAAGAAGAAGATAGTAGACGCTACCTAGCACCGAG    1140
sequencing_primer_2       AGGAGTACAAGAACTTCGAAAACGACAAGAAGAAGATAGTAGACGCTAACATAGCAACAG    677

K13_C580Y_sequence        AGACAATGATAGACATAAACGTAGGAGGTGCAATATTCGAGACTTCAAGGCACACTTTGA    1105
sequencing_primer_2       AGACAATGATAGACATAAACGTAGGAGGTGCAATATTCGAGACTTCAAGGCACACTTTGA    737

K13_C580Y_sequence        CTCAGCAGAAGGACTCTTTCATTGAAAAGTTGTTGTCAGGTAGGCACCACGTTACAAGGG    1165
sequencing_primer_2       CTCAGCAGAAGGACTCTTTCATTGAAAAGTTGTTGTCAGGTAGGCACCACGTTACAAGGG    797

K13_C580Y_sequence        ACAAGCAGGGTAGGATTTTTTTGGACAGAGACTCAGAATTGTTCAGGATAATTTTAAATT    1225
```

```
sequencing_primer_2      ACAAGCAGGGTAGGATTTTTTTGGACAGAGACTCAGAATTGTTCAGGATAATTTTAAATT    857

K13_C580Y_sequence       TCTTGAGGAACCCCTTGACAATTCCTATTCCTAAGGACTTGTCTGAGTCAGAGGCTTTAT    1285
sequencing_primer_2      TCTTGAGGAACCCCTTGACAATTCCTATTCCTAAGGACTTGTCTGAGTCAGAGGCTTTAT    917

K13_C580Y_sequence       TAAAGGAGGCTGAGTTCTACGGAATAAAGTTCTTGCCTTTTCCTTTGGTTTTCTGCATTG    1345
sequencing_primer_2      TAAAGGAGGCTGAGTTCTACGGAATAAAGTTCTTGCCTTTTCCTTTGGTTTTCTGCATTG    977

K13_C580Y_sequence       GAGGTTTCGACGGAGTTGAGTACTTGAACAGTATGGAGTTGTTGGACATATCACAGCAGT    1405
sequencing_primer_2      GAGGTTTCGACGGAGTTGAGTACTTGAACAGTATGGAGTTGTTGGACATATCACAGCAGT    1037

K13_C580Y_sequence       GTTGGCGAATGTGCACTCCAATGTCAACTAAGAAGGCATACTTCGGTTCTGCAGTTTTAA    1465
sequencing_primer_2      GTTGGCGAATGTGCACTCCAATGTCAACTAAGAAGGCATACTTCGGTTCTGCAGTTTTAA    1097

K13_C580Y_sequence       ACAACTTTTTGTATGTATTCGGAGGAAACAATTACGACTACAAAGCATTGTTCGAGACAG    1525
sequencing_primer_2      ACAACTTTTTGTATGTATTCGGAGGAAACAATTACGACTACAAAGCATTGTTCGAGACAG    1157

K13_C580Y_sequence       AGGTATACGACAGGTTGAGGGACGTTTGGTACGTAAGTTCAAACTTGAACATTCCAAGGA    1585
sequencing_primer_2      AGGTATACGACAGGTTGAGGGACGTTTGGTACGTAAGTTCAAACTTGAACATTCCAAGGA    1217

K13_C580Y_sequence       GGAACAACTGCGGAGTAACCTCTAACGGAAGGATATACTGCATAGGAGGTTACGACGGAT    1645
sequencing_primer_2      GGAACAACTGCGGAGTAACCTCTAACGGAAGGATATACTGCATAGGAGGTTACGACGGAT    1277
sequencing_primer_3      ------ACTGCGGAGTAACCTCTAACGGAAGGATATACTGCATAGGAGGTTACGACGGAT    72

K13_C580Y_sequence       CATCAATAATTCCCAACGTTGAGGCTTACGACCACAGAATGAAGGCTTGGGTTGAAGTAG    1705
sequencing_primer_3      CATCAATAATTCCCAACGTTGAGGCTTACGACCACAGAATGAAGGCTTGGGTTGAAGTAG    132

K13_C580Y_sequence       CTCCATTAAACACTCCAAGGTCTAGTGCAATGTATGTAGCATTCGACAACAAGATATACG    1765
sequencing_primer_3      CTCCATTAAACACTCCAAGGTCTAGTGCAATGTATGTAGCATTCGACAACAAGATATACG    192

K13_C580Y_sequence       TGATAGGAGGTACAAACGGAGAAAGGTTGAACTCAATAGAGGTTTACGAGGAGAAGATGA    1825
sequencing_primer_3      TGATAGGAGGTACAAACGGAGAAAGGTTGAACTCAATAGAGGTTTACGAGGAGAAGATGA    252

K13_C580Y_sequence       ACAAGTGGGAGCAGTTCCCTTACGCATTGTTGGAGGCAAGGTCAAGTGGTGCTGCATTCA    1885
sequencing_primer_3      ACAAGTGGGAGCAGTTCCCTTACGCATTGTTGGAGGCAAGGTCAAGTGGTGCTGCATTCA    312

K13_C580Y_sequence       ACTATTTAAACCAGATTTACGTAGTAGGTGGAATAGACAACGAGCACAATATTTTGGACT    1945
sequencing_primer_3      ACTATTTAAACCAGATTTACGTAGTAGGTGGAATAGACAACGAGCACAATATTTTGGACT    372

K13_C580Y_sequence       CTGTAGAGCAGTACCAGCCTTTCAACAAGAGGTGGCAGTTCCTTAACGGAGTTCCTGAAA    2005
sequencing_primer_3      CTGTAGAGCAGTACCAGCCTTTCAACAAGAGGTGGCAGTTCCTTAACGGAGTTCCTGAAA    432

K13_C580Y_sequence       AGAAGATGAACTTCGGTGCAGCTACTTTAAGTGACTCATACATTATAACTGGTGGTGAGA    2065
sequencing_primer_3      AGAAGATGAACTTCGGTGCAGCTACTTTAAGTGACTCATACATTATAACTGGTGGTGAGA    492

K13_C580Y_sequence       ACGGGGAGGTACTTAACTCTTGCCACTTTTTCTCACCTGACACTAACGAGTGGCAATTAG    2125
sequencing_primer_3      ACGGGGAGGTACTTAACTCTTGCCACTTTTTCTCACCTGACACTAACGAGTGGCAATTAG    552

K13_C580Y_sequence       GACCTTCATTGTTGGTACCAAGGTTCGGACATTCAGTATTGATTGCTAACATTTGA----    2181
sequencing_primer_3      GACCTTCATTGTTGGTACCAAGGTTCGGACATTCAGTATTGATTGCTAACATTTGAACCG    612
```

**Fig. S2.**

**Sanger sequencing results of the k13 gene in K13$^{C580Y}$-29$^{th}$ parasites.** CLUSTAL O(1.2.4) multiple sequence alignment.

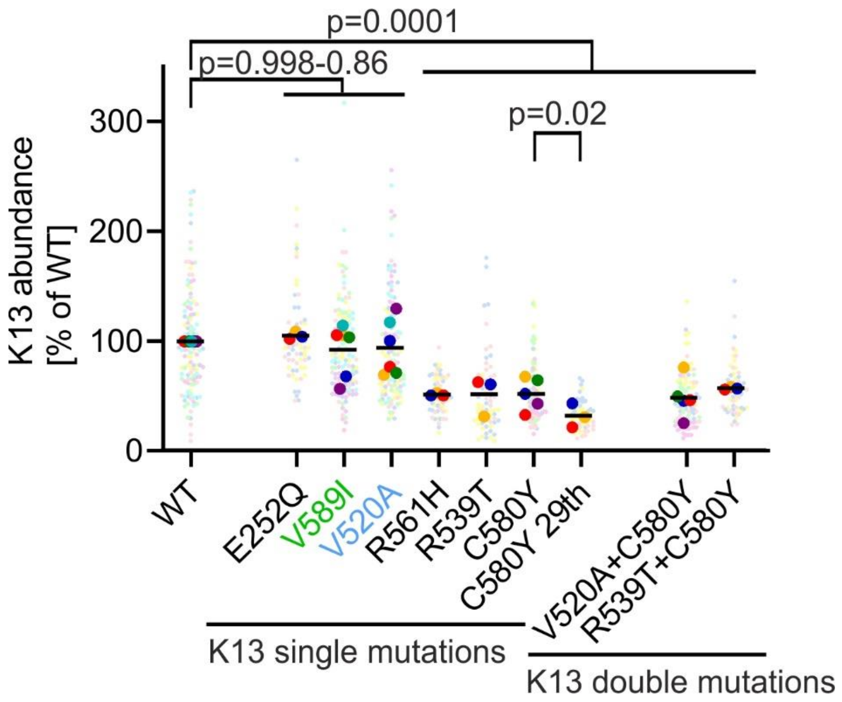


**Fig. S3.**

**K13 quantification with individual data point used for statistical analysis.** K13-abundance measured by total GFP fluorescence intensity of parasites with the indicated K13 expressed from the endogenous locus and fused to GFP, normalized to the fluorescence in parasites with the identically modified endogenous locus but with a WT GFP-K13. Small dots represent measured values. Large dots of the same color as small dots represent mean of the respective measured values and each large dot represents one biological replicate. Same data as in Figure 3A. P-values derive from comparing values from each measured cell by one-way ANOVA.

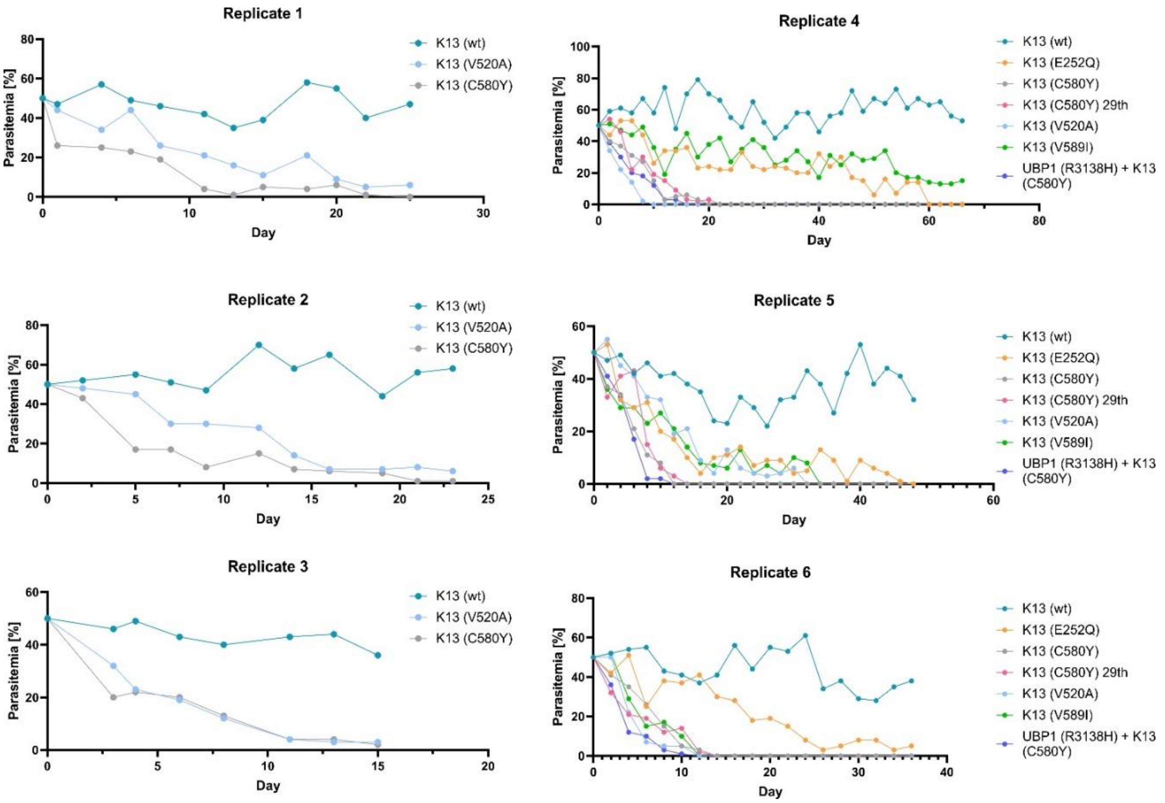


**Fig. S4.**

**Growth curves of fitness assays.** Raw curves of six independent fitness assays performed in RPMI complete medium.

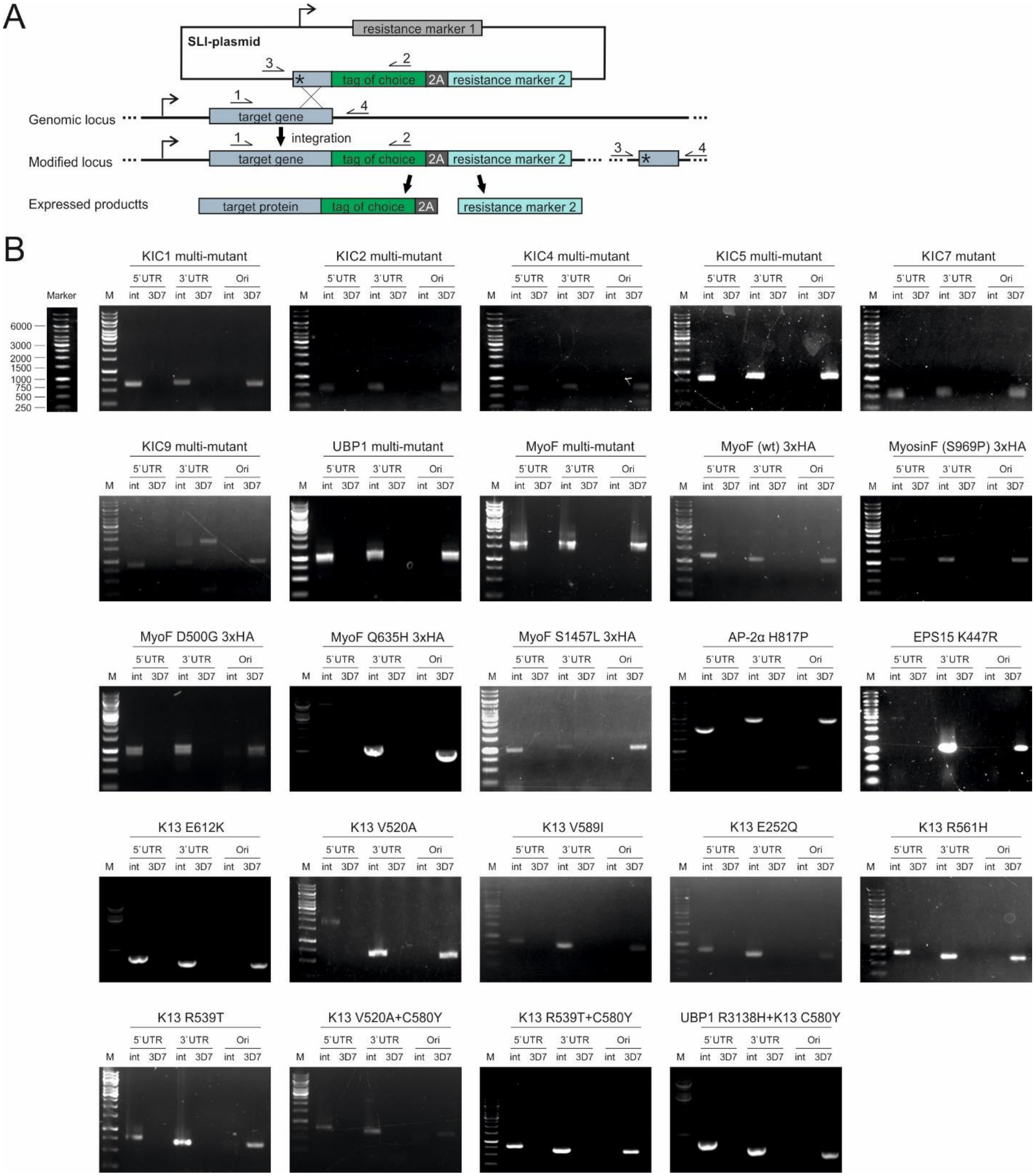


**Fig. S5.**

**Confirmation of correct integration of genome-modified parasites. (A)** Schematic of selection-linked integration (SLI). **(B)** The correct integration was confirmed by generating PCR products from gDNA of indicated parasites lines, subsequently separated on agarose gels. Primers generate products across the 5' and 3' integration junctions (labelled 5' and 3') and the unmodified locus (ori), which is not present in SLI-edited lines (int) but in the parental parasites 3D7. Wild type (WT). Marker (M) fragment length is indicated in base pairs.

**Table S1.**
**List of all SNPs collected for multi-mutants pools and for individual testing of the candidate genes indicated.** Asterisk (*), indicates SNP that were tested individually and not in the multi-mutant since they were detected after SLI plasmids of the multi-mutants were prepared for transfection or were located in the N-terminal region of the gene which would have complicated their inclusion due to the large size of this gene, and were therefore tested individually.

| Candidate | Mutations |
|---|---|
| *Pf*KIC1 (PF3D7_0606000) | D483H, D535G, D862V, I873M, K231E, K676N, K883T, N179K, N633Y, N634Y, N688Y, N971S, Q1042K, S394N, S429P, T679A, V543F |
| *Pf*KIC2 (PF3D7_1227700) | E207Q, E343K, G167D, K321N, K651M, N449T, Y472F |
| *Pf*KIC4 (PF3D7_1246300) | D120N, I121V, M375V, N123S, N405S, N624I, N661H, P170S, P621A, T888A, V619A, Y179C |
| *Pf*KIC5 (PF3D7_1138700) | A471T, D169N, E1055G, E1354D, E1646G, E497Q, F570L, G1417D, G264D, G727E, H1198Y, H138Y, K1411R, K258R, K384I, K751Q, K985N, L1066P, M324V, M975I, N1268H, N525S, P516S, Q110E, Q1127K, Q1421K, R1040K, S1044R, S1721A, T1381A, V348I, V72I, V836I |
| *Pf*KIC7 (PF3D7_0813000) | I130V, N434S, N596K, P318S, S576N |
| *Pf*KIC9 (PF3D7_1442400) | D1345N, D1879V, D2018E, D388E, D734Y, E1861K, G674V, G984R, H1710R, I1280L, I1688V, I429S, I960V, K1039N, K1626N, K589R, K837I, L279I, M1507I, M618T, N1402K, N286H, N927H, P249Q, Q1045E, S1549N, S282N, S341N, S753T, T990R, T996I, V1586I, V2072M, Y293N |
| *Pf*MyosinF (PF3D7_1329100) | D500G*, Q635H*, S969P*, S1457L*, C1196S, G873D, H1587R, I2068T, L1165V, M1872L, N1615K, V1193E |
| *Pf*UBP1 (PF3D7_0104300) | D2704E, D2925N, G2810C, K3013Q, M2618V, N2165I, N2669S, Q2355E, Y2530H |
| *Pf*Eps15 | K447R* |
| *Pf*AP-2α | H817P* |

**Table S2.**
**Parasite lines used in this study.**

| Parasite line | Tag | Source |
|---|---|---|
| 3D7 | none | Walliker et al., 1987 |
| KIC1 WT | Sandwich | Birnbaum et al., 2020 |
| KIC1 Mut | GFP | This study |
| KIC2 WT | Sandwich | Birnbaum et al., 2020 |
| KIC2 Mut | GFP | This study |
| KIC4 WT | Sandwich | Birnbaum et al., 2020 |
| KIC4 Mut | GFP | This study |
| KIC5 WT | Sandwich | Birnbaum et al., 2020 |
| KIC5 Mut | GFP | This study |
| KIC7 WT | Sandwich | Birnbaum et al., 2020 |
| KIC7 Mut | GFP | This study |
| KIC9 WT | Sandwich | Birnbaum et al., 2020 |
| KIC9 Mut | GFP | This study |
| UBP1 WT | 2xFKBP-GFP | Birnbaum et al., 2020 |
| UBP1 Mut | GFP | This study |
| MyoF Mut | GFP | This study |
| MyoF WT 3xHA | 3xHA | This study |
| $MyoF^{S969P}$ | 3xHA | This study |
| $MyoF^{D500G}$ | 3xHA | This study |
| $MyoF^{Q635H}$ | 3xHA | This study |
| $MyoF^{S1457L}$ | 3xHA | This study |
| $AP\text{-}2\alpha^{H817P}$ | 3xHA | This study |
| $EPS15^{K447R}$ | GFP | This study |
| K13 WT | N-GFP-2xFKBP | Birnbaum et al., 2017 |
| $K13^{E612K}$ | N-GFP-2xFKBP | This study |
| $K13^{V520A}$ | N-GFP-2xFKBP | This study |
| $K13^{V589I}$ | N-GFP-2xFKBP | This study |
| $K13^{E252Q}$ | N-GFP-2xFKBP | This study |
| $K13^{C580Y}$ | N-GFP-2xFKBP | Birnbaum et al., 2017 |
| $K13^{R561H}$ | N-GFP-2xFKBP | This study |
| $K13^{R539T}$ | N-GFP-2xFKBP | This study |
| $K13^{V520A+C580Y}$ | N-GFP-2xFKBP | This study |
| $K13^{R539T+C580Y}$ | N-GFP-2xFKBP | This study |
| $K13^{C580Y}$-29$^{th}$ | N-GFP-2xFKBP | This study |
| K13 1xNLS | N-GFP-2xFKBP + mCherry | Birnbaum et al., 2020 |
| KIC7 1xNLS | N-GFP-2xFKBP + mCherry | Birnbaum et al., 2020 |
| $UBP1^{R3138H}$ | 3xHA | Birnbaum et al., 2020 |
| $UBP1^{R3138H}$ +$K13^{C580Y}$ | 3xHA + N-GFP-2xFKBP | This study |

**Dataset S1 (separate file).**
List of *k13* V520A, V589I and E612K mutations found in Africa. Data were extracted from WWARN Artemisinin Molecular Surveyor database.

**Dataset S2 (separate file).**
Source data for Figure 4E. Malaria incidence at place and time of *k13* resistance mutation detection.

**Dataset S3 (separate file).**
Primers used in this study.
